# DLGAP5 promotes vascular cell proliferation and remodeling in pulmonary arterial hypertension

**DOI:** 10.64898/2026.09.29.755207

**Authors:** Tapan Dey, Iryna Zhyvylo, Dmitry Goncharov, Derek Lin, John R. Greenland, Paul Wolters, Delphine Gomez, Lifeng Jiang, Elena A. Goncharova

**Author notes:** **Correspondence to:** Elena A. Goncharova, PhD, GBSF Rm 6523, 451 Health Sciences Drive, Davis, CA 95616.

## Abstract

**Background:** Pulmonary arterial hypertension (PAH) manifests by pulmonary vascular remodeling, caused by hyper-proliferation and apoptosis resistance of resident PA cells, elevated right ventricular (RV) afterload, and death of RV failure. The goal of this study is to determine the role and mechanisms of action of mitotic spindle-associated microtubule regulator DLG associated protein 5 (DLGAP5) in pulmonary vascular remodeling in PAH.

**Methods:** DLGAP5 expression was assessed in human PAH transcriptomic datasets, PAH patients’ lung tissues, pulmonary artery smooth muscle cells (PASMCs), and pulmonary artery adventitial fibroblasts (PAAFs). Loss-of-function, molecular, functional, RNAseq, and network analyses were performed. Smooth muscle (SM)-specific DLGAP5 knockdown using adeno-associated virus serotype 6 (AAV6) in mice with SU5416/hypoxia-induced PH was performed.

**Results:** DLGAP5 was overaccumulated in small muscular PAs, PASMCs, and PAAFs from PAH lungs. siRNA DLGAP5 down-regulated cyclin A2, suppressed hyper-proliferation, increased pro-apoptotic BIM, and induced apoptosis in human PAH PASMCs and PAAFs. DEP domain containing 1 (DEPDC1) was identified and validated as a downstream effector of DLGAP5 in PAH PASMCs responsible for hyper-proliferation and apoptosis resistance of human PAH PASMCs. SM-specific DLGAP5 knockdown with AAV6-*enTagln*-shDLGAP5 reversed DEPDC1 overaccumulation in small muscular PAs, reduced PA medial thickness, RV systolic pressure, mean PA pressure, and RV hypertrophy in mice with SuHx-induced PH.

**Conclusions:** DLGAP5 supports proliferation and survival of human PAH PASMCs and PAAFs through up-regulation of cyclin A2, DEPDC1, and suppression of BIM, and promotes PA remodeling and PH in mice. DLGAP5 could serve as a potential molecular target to attenuate pulmonary vascular remodeling in PAH.

**Graphical abstract:** 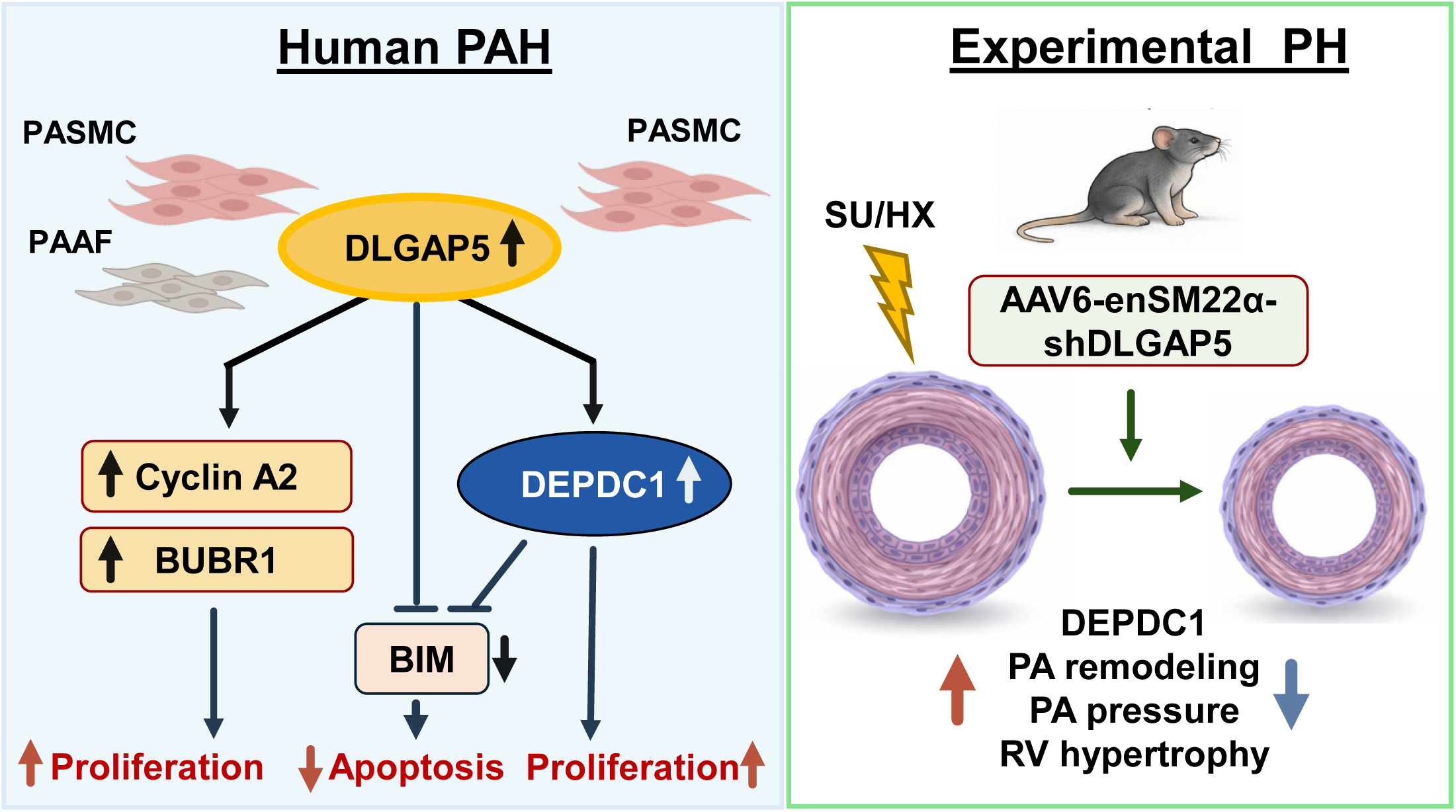

## INTRODUCTION

Pulmonary arterial hypertension (PAH) is a progressive cardiovascular disease characterized by sustained elevation of pulmonary artery (PA) pressure, increased pulmonary vascular resistance, and progressive right ventricular (RV) overload, ultimately leading to RV failure and premature mortality^1,2^. A defining pathological feature of PAH is structural remodeling of distal pulmonary arteries (PAs), driven in part by pathological accumulation and phenotypic reprogramming of pulmonary artery smooth muscle cells (PASMCs) and pulmonary artery adventitial fibroblasts (PAAFs)^3–7^. These vascular mesenchymal cells acquire a hyperproliferative apoptosis-resistant phenotype that contributes to PA remodeling, persistent elevation of PA pressure and pulmonary vascular resistance^3–7^. Although most established PAH therapies improve symptoms and hemodynamics by targeting vasoactive pathways, only one FDA-approved drug, Sotatercept, targets PA remodeling in PAH^8^, and new remodeling-focused strategies are urgently needed^1,9,10^.

Proliferative and apoptosis-resistant phenotype of PAH pulmonary vascular cells is supported by multiple factors, including dysregulated growth factor signaling, metabolic reprogramming, inflammation, and extracellular matrix remodeling^11–18^. Sustained hyper-proliferation requires up-regulation of cell cycle progression and mitotic regulatory programs, including spindle assembly, chromosome alignment, and checkpoint control^19–23^. Perturbation of spindle assembly or mitotic checkpoint signaling can result in mitotic arrest, chromosome mis-segregation, mitotic catastrophe, inflammatory signaling, or apoptosis^19,20,23,24^. In cancer biology, mitotic regulators are increasingly recognized not only as mediators of chromosome segregation, but also as determinants of cell survival and disease progression^24^. Several studies have highlighted the importance of mitotic regulation in PAH. Dynamin-related protein-1 (DRP1), microtubule motor protein Kinesin Family Member 23 (KIF23), and mitotic checkpoint protein BUB3 are identified as important regulators of PASMC proliferation, PA remodeling, and PH; and genes involved in the mitotic cell cycle and microtubule cytoskeleton organization are enriched in human PAH lungs^17,25–29^.

DLG associated protein 5 (DLGAP5), also known as hepatoma upregulated protein (HURP), is a microtubule-associated protein that localizes to the mitotic spindle and supports spindle dynamics, kinetochore capture, Ran-dependent spindle formation, and mitotic progression^30,31^. DLGAP5 expression is closely linked to cell proliferative state. DLGAP5 is elevated in several malignancies, promoting cell cycle progression, proliferation, invasion, and survival^32–35^. The role of DLGAP5 in pulmonary vascular remodeling and pulmonary hypertension (PH) has not been defined. Given the dependence of PAH vascular cells on sustained proliferation and apoptosis resistance, we hypothesized that aberrant DLGAP5 upregulation may contribute to pulmonary vascular remodeling by supporting cell-cycle progression and mitotic regulatory programs.

To test this hypothesis, we examined DLGAP5 expression in human PAH lung tissue and patient-derived PASMCs, PAAFs, and PA endothelial cells (PAECs), and evaluated the functional consequences of DLGAP5 loss in PAH pulmonary vasculature. We further investigated DLGAP5-regulated mitotic programs and downstream effectors and assessed consequences of smooth muscle (SM)-targeted DLGAP5 knockdown in experimental PH *in vivo.* Through these complementary approaches, we sought to define whether DLGAP5 represents a disease-relevant molecular target to reverse pulmonary vascular remodeling and overall PH.

## MATERIALS AND METHODS

Detailed methods are provided in the Online Supplement.

### Human tissues and cells

Human lung tissues and early-passage (3–8) primary human PASMCs, PAAFs, and PAECs were obtained from the Pulmonary Hypertension Breakthrough Initiative (PHBI), the University of California, Davis Lung Center Pulmonary Vascular Disease Program biobank (tissue acquired from the University of Pittsburgh Medical Center Lung Transplant program and Pulmonary Division Tissue Donation program, Institutional Review Board (IRB) approvals #PRO14010265, #PRO13050218, and Committee for Oversight of Research Involving Decedents approvals #451, #458), and the University of California San Francisco Lung Transplant Program (IRB protocol # 13-10738). Human subject characteristics are summarized in Table S1. Isolation, characterization, and maintenance of PASMCs, PAAFs, and PAECs were performed according to established PHBI protocols^14,16,17,36^. Experiments were performed using primary cells from independent human subjects as biological replicates.

### Animal studies

All animal procedures were performed under protocols approved by the Institutional Animal Care and Use Committee of the University of California, Davis. Twelve-week-old male C57BL/6J mice were purchased from The Jackson Laboratory (Bar Harbor, ME). Animals were randomized to experimental groups, blinded hemodynamic and histological analyses were performed. Mice were exposed to normobaric hypoxia (10%O₂) for 7 weeks and received SU5416 (20 mg/kg, s.c.) on days 1, 7, and 14 of experiment. On day 1, mice received a single tail-vein injection of AAV6 encoding an *enTagln*-driven DLGAP5-targeting shRNA or a matched scrambled control shRNA at a dose of 1×10¹¹ viral genomes (vg)/mouse. Age-matched male C57BL/6J mice maintained under normoxic conditions were used as controls. At the end of the 7 week hypoxia exposure, terminal hemodynamic analysis was performed as described before^14,15,17,18^, mean pulmonary artery pressure (mPAP) was calculated as 0.61 x sRVP + 2 (in mmHg)^37^, PA Medial thickness (PA MT) was quantified using the VMI calculator^38^, Fulton index was calculated as the RV/(LV+septum) weight ratio^14,17,18^.

### Transcriptomic analyses

Publicly available human PAH transcriptomic datasets GSE113439^39^, GSE144274^40^, GSE193776^41^ and GSE126262^42^ were analyzed to evaluate DLGAP5 expression in whole lung tissue and isolated PASMCs, PAAFs, and PAECs, respectively. Bulk RNA sequencing was performed in PASMCs from three patients with PAH transfected with control siRNA (ON-TARGETplus nontargeting siRNA #1) or siRNA DLGAP5. RNAseq and pathway enrichment analyses were performed by Genenwiz® Azenta Life Sciences (South Plainfield, NJ).

### Immunoblot, immunohistochemistry, immunocytochemistry, transfection and functional analyses

Immunoblotting, immunohistochemistry, immunocytochemistry, cell transfection, cell proliferation, and apoptosis analyses were performed as described before in^14,15,17,18^.

### Statistical analysis

Statistical analysis was performed using StatView and GraphPad Prism software. For datasets with n<6/group, or datasets with n≥6/group without normal distribution, differences between 2 groups were assessed using Mann-Whitney U test. Normally distributed datasets with n≥6 per group were analyzed using Student’s t test for comparisons between 2 groups or one-way ANOVA followed by Dunnett’s post hoc multiple comparisons for ≥3 groups. Statistical significance was defined as p<0.05.

## RESULTS

### DLGAP5 is upregulated in pulmonary vascular mesenchymal cells in PAH lungs

To determine the status of DLGAP5 in PAH, we analyzed publicly available datasets GSE113439^39^, GSE144274^40^, and GSE193776^41^. We found that normalized DLGAP5 expression was significantly higher in human PAH lungs, PASMCs and PAAFs compared to respective controls (Figure 1A). Immunohistochemical analysis demonstrated that DLGAP5 is overaccumulated in smooth muscle α-actin (SMA)-positive areas of small muscular PAs in PAH lungs (Figure 1B). Immunoblot analysis of distal early-passage PASMCs and PAAFs showed that DLGAP5 protein levels are significantly higher in human PAH PASMCs and PAAFs compared to controls (Figure 1C-F). In contrast, we detected no appreciable differences in DLGAP5 mRNA (GSE126262)^42^ or protein levels (Figure S1) between control and PAH PAECs. Taken together, these findings indicate that DLGAP5 is upregulated in PASMCs and PAAFs, but not PAECs, in human PAH lungs.

**Figure 1.**
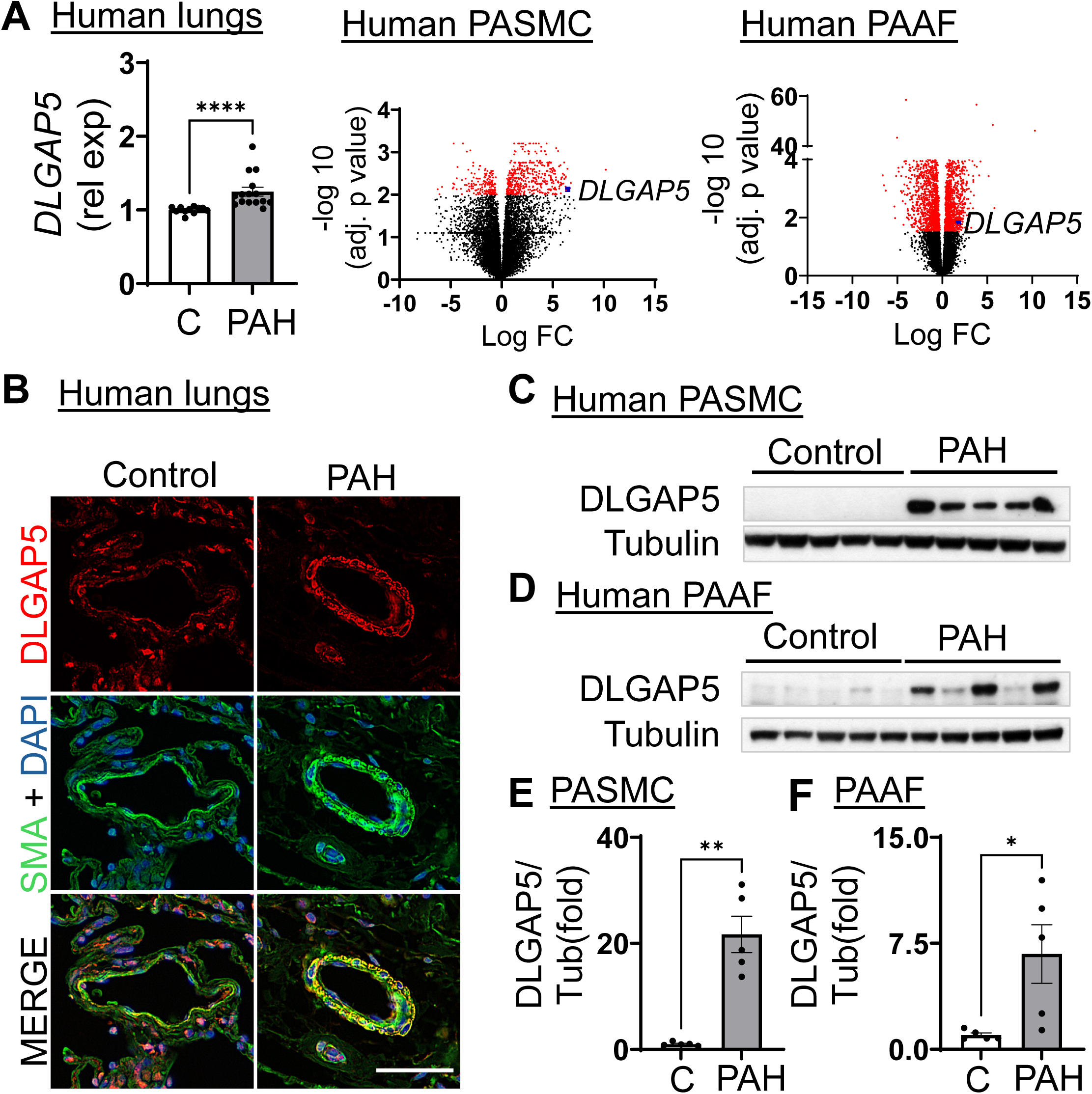
DLGAP5 is upregulated in human PAH pulmonary arteries and isolated pulmonary vascular cells. **A**: DLGAP5 transcript level in human control and PAH lungs; GSE113439 (control, n=11; PAH, n=15) (left); volcano plots of differentially expressed genes in PASMCs (GSE144274) (middle) and PAAFs (GSE193776) (right) from control and PAH subjects (n=4 subjects/group). Significantly differentially expressed genes (DEGs) are shown in red, DLGAP5 is highlighted. **B**: Representative images of human control and PAH lungs stained for DLGAP5 (red), smooth muscle α-actin (SMA; green), and DAPI (blue) from 5 subjects/group. Bar=50 μm. **C-F**: Immunoblot analysis of indicated proteins in PASMCs (C, E) and PAAFs (D, F) from control and PAH subjects. Data are means±SEM; n=5 subjects/group; each point represents an individual subject. *p<0.05; **p<0.01, ****p<0.0001 by Mann-Whitney U test.

### DLGAP5 supports proliferation, survival, and mitotic regulatory proteins in PAH pulmonary vascular cells

To determine the functional significance of DLGAP5 in human PAH PASMCs and PAAFs, we depleted DLGAP5 using specific siRNA and examined whether its depletion affects proteins that regulate cell cycle progression, mitotic checkpoint, and cell survival. We found that DLGAP5 depletion reduced protein levels of budding uninhibited by benzimidazole-related 1 (BUBR1), a key component of the spindle-assembly checkpoint that is required for proper chromosome segregation during mitosis^43,44^, and down-regulated cyclin A2, an important regulator of cell proliferation that is upregulated in experimental PH^45–47^. Furthermore, DLGAP5 loss significantly increased protein levels of proapoptotic BH3-only protein BIM, deficiency of which is responsible for apoptosis resistance of human PAH PASMCs^14^ (Figure 2A, D). These molecular changes were accompanied by significant decrease in cell proliferation and induction of apoptosis compared to cells transfected with control siRNA (Figure 2B, C, E, F). These data show that DLGAP5 supports overaccumulation of mitotic regulatory proteins cyclin A2 and BUBR1, deficiency of pro-apoptotic BIM, hyper-proliferation and apoptosis resistance of human PAH PASMCs and PAAFs.

**Figure 2.**
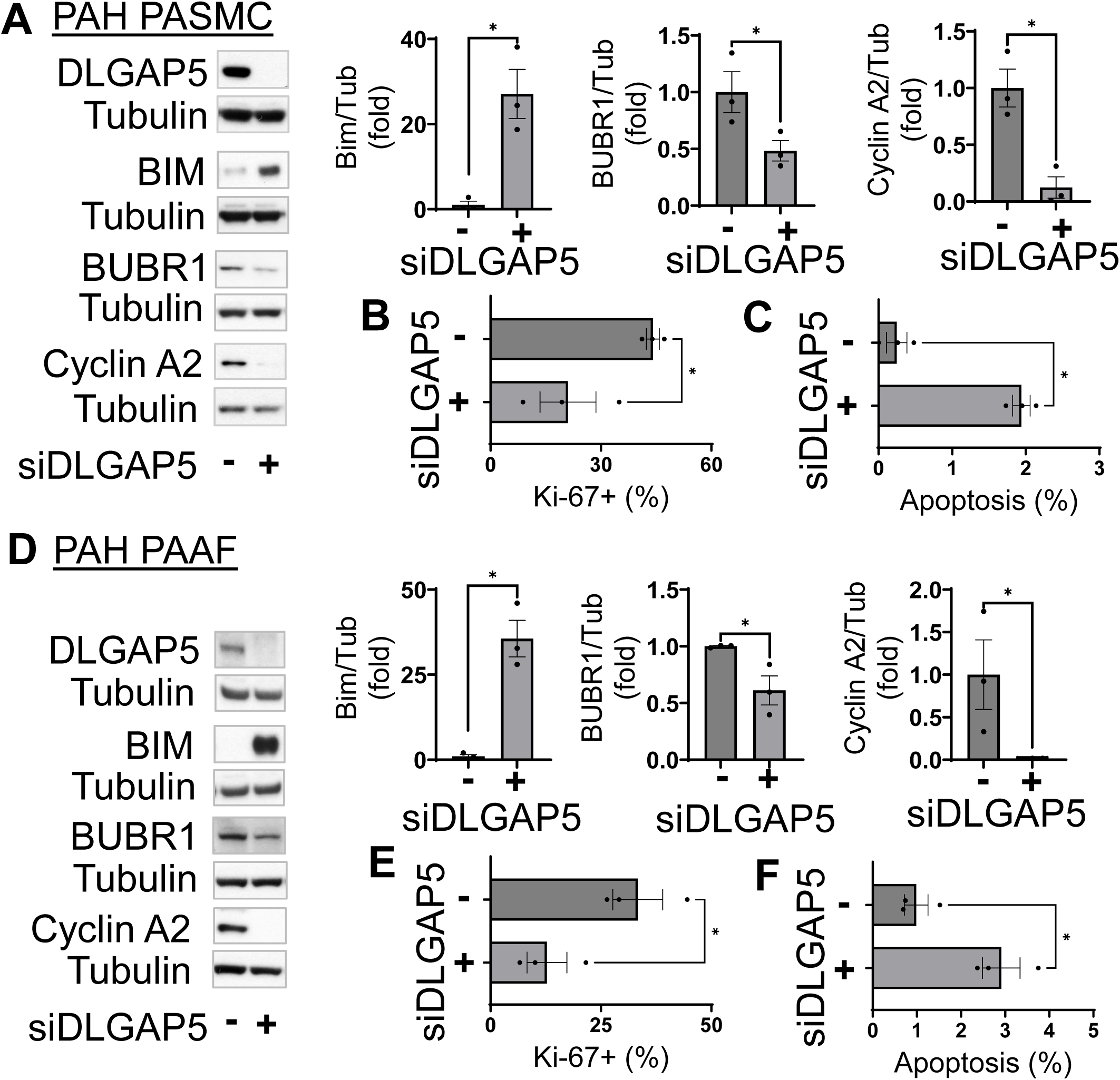
DLGAP5 depletion suppresses mitotic regulators, reduces proliferation, and induces apoptosis in PAH PASMCs and PAAFs. PAH PASMCs and PAAFs were transfected with control scrambled siRNA (-) or siRNA DLGAP5 (siDLGAP5; +) for 48 hours. Then, Immunoblot analysis of indicated proteins ( **A, D**), proliferation (Ki-67) (**B, E**), and apoptosis analyses (*In Situ* Cell Death Detection Kit) (**C, F**) were performed. Data are means±SEM; images are representative from n=3 subjects/group. *p<0.05 by Mann-Whitney U test.

### DLGAP5 modulates human PAH PASMC proliferation and survival via DEPDC1

To define transcriptional programs regulated by DLGAP5, we performed bulk RNAseq of human PAH PASMCs transfected by siRNA DLGAP5 and control siRNA. It identified 100 differentially expressed genes, including 80 up- and 20 downregulated (Figure 3A). Gene ontology analysis revealed enrichment of biological processes associated with nucleosome and chromatin organization, DNA packaging, mitotic spindle assembly and elongation, and spindle midzone organization (Figure 3A).

**Figure 3.**
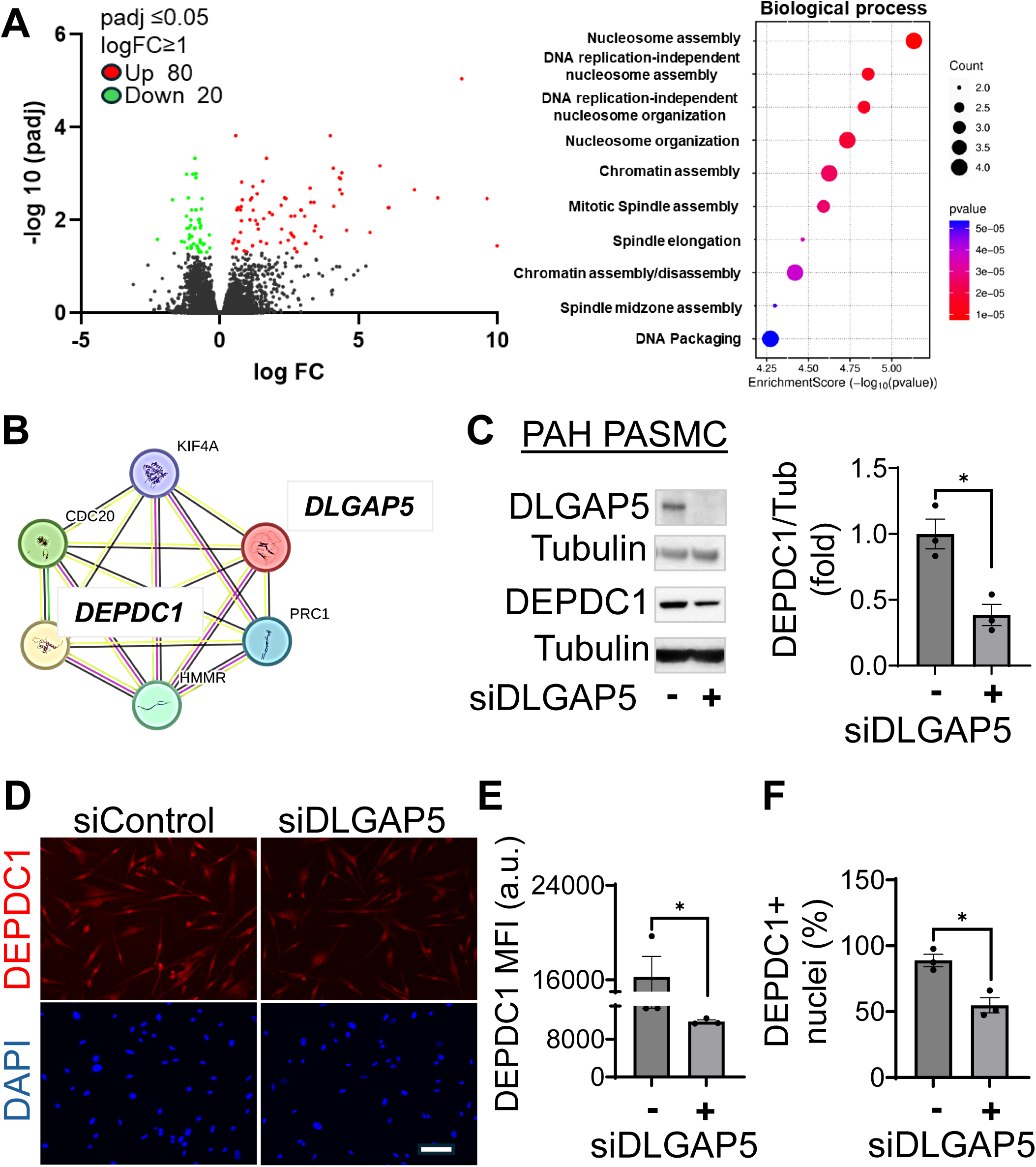
DLGAP5 supports DEPDC1 expression and nuclear accumulation in human PAH PASMCs. **A:** human PAH PASMCs were transfected with DLGAP5-targeting siRNA (siDLGAP5) or scrambled control siRNA for 48 hours, and bulk RNA sequencing was performed. Left: Volcano plot of differentially expressed genes (DEGs). Significantly upregulated and downregulated DEGs are shown in red and green, respectively (adjusted P≤0.05, log2FC≥1; n=3 subjects/group). Right: Gene Ontology biological processes enriched among significantly downregulated DEGs. Dot size represents the number of associated genes, color represents statistical significance. **B**: STRING protein-protein association network constructed from significantly downregulated DEGs. Nodes represent the encoded proteins, edges represent known or predicted functional associations. **C**: Immunoblot analysis of human PAH PASMCs transfected with scrambled control siRNA or siDLGAP5 for 48 hours. **D-F**: Immunocytochemical analysis of human PAH PASMCs transfected with siControl or siDLGAP5 for 48 hours. Red - DEPDC1, blue - DAPI Representative images from n=3 subjects/group (**D**), background-corrected nuclear DEPDC1 mean fluorescence intensity (MFI) (**E**), and the percentage of cells with nuclear DEPDC1 signal (**F**) were shown. Nuclear DEPDC1 MFI was measured within DAPI-defined nuclear regions of interest. DEPDC1-positive nuclei were defined using a background-corrected MFI threshold of >11,000 arbitrary units, applied uniformly across all samples. Bar=100μm. Data are means±SEM, 3 subjects/group, *p<0.05 by Mann-Whitney U test.

Protein-protein interaction analysis of genes downregulated by DLGAP5 knockout using Search Tool for the Retrieval of Interacting Genes/Proteins (STRING) identified an interconnected module containing DLGAP5, DEP domain containing 1 (DEPDC1), kinesin family member 4A (KIF4A), protein regulator of cytokinesis 1 (PRC1), cell division cycle protein 20 (CDC20), and hyaluronan mediated motility receptor (HMMR) (Figure 3B). Supporting the relevance of our findings to human disease, CDC20 was previously linked to hyper-proliferation of PASMCs and PAAFs and PAH pathogenesis^48^. Out of other four genes, DEPDC1 was of particular interest because of its established association with proliferative programs^49–51^.

To validate RNAseq-based data, we silenced DLGAP5 in PAH PASMCs using specific siRNA and evaluated DEPDC1 protein levels and cellular localization. DLGAP5 depletion significantly reduced DEPDC1 protein levels and decreased % of nuclear DEPDC1 and % of cells with nuclear DEPDC1 accumulation compared to the cells transfected with control siRNA (Figure 3C-E), showing that DLGAP5 supports DEPDC1 nuclear overaccumulation in human PAH PASMCs.

To evaluate an applicability of our findings to human PAH, we next tested the status of DEPDC1 in human PAH pulmonary vascular cells. Analysis of publicly available datasets demonstrated that DEPDC1 expression is increased in lungs, PASMCs, and PAAFs from PAH patients compared to non-diseased controls (Figure 4A). Supporting RNAseq-based data, DEPDC1 protein levels were significantly higher in human PAH PASMCs (Figure 4B). Interestingly, we found no differences in DEPDC1 protein content between control and PAH PAAFs (Figure 4C), suggestive of PASMC-specific DLGAP5 action. Transfection of human PAH PASMCs with siRNA DEPDC1 significantly increased BIM protein levels (Figure 4D), reduced percentage of Ki-67-positive cells (Figure 4E) and induced significant apoptosis (Figure 4F) compared to cells transfected with control siRNA. Interestingly, in contrast to siDLGAP5, siRNA DEPDC1 did not affect BUBR1 and cyclin A2 protein levels (Figure S2), suggesting that DLGAP5 modulates DEPDC1 in parallel to BUBR1 and cyclin A2. Collectively, these data demonstrate that DEPDC1 acts downstream of DLGAP5 to promote Bim deficiency, hyper-proliferation and survival of human PAH PASMCs.

**Figure 4.**
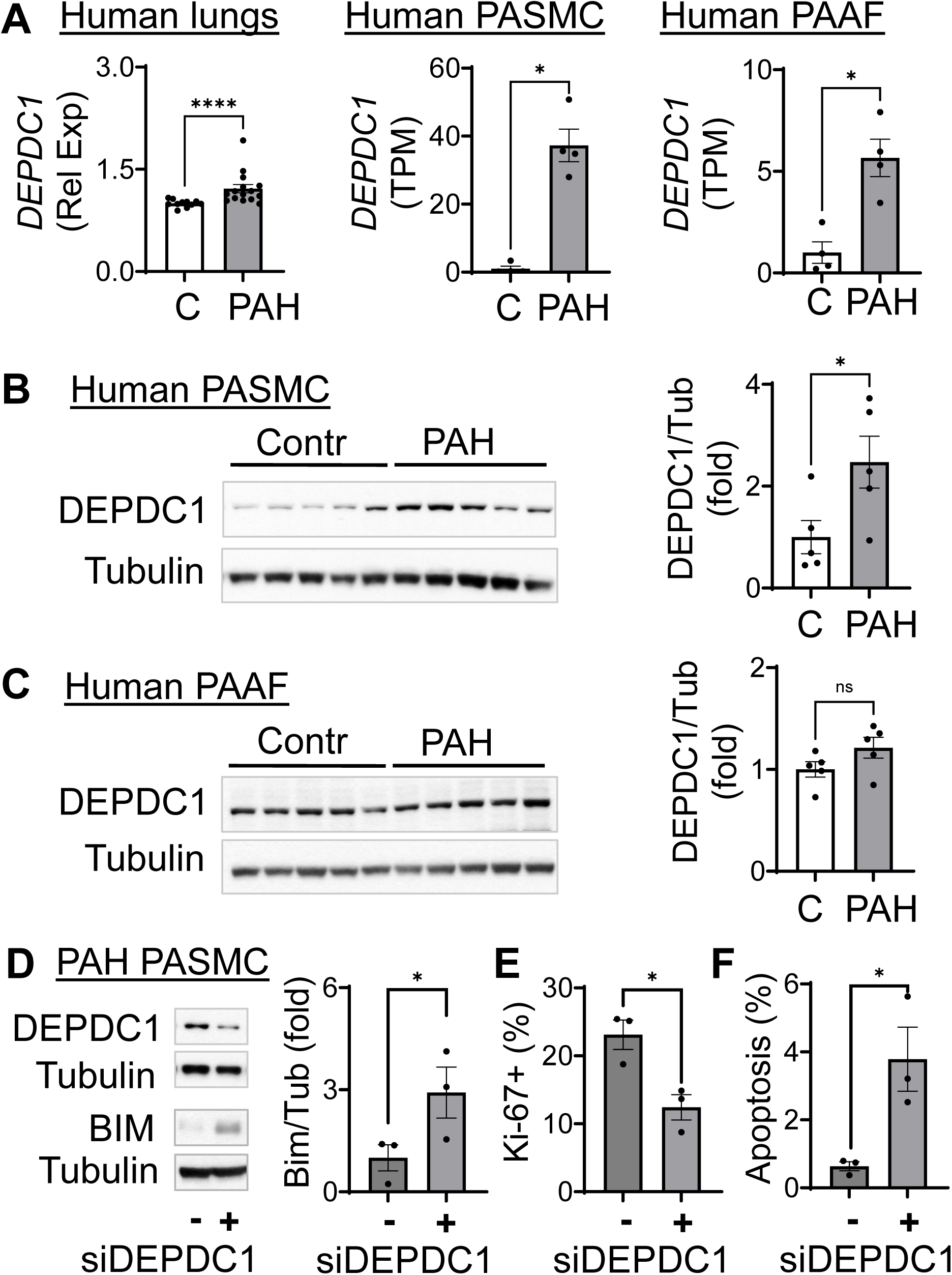
DEPDC1 depletion suppresses proliferation and promotes apoptosis in human PAH PASMCs. **A:** *DEPDC1* expression in human control and PAH lungs (GSE113439) (left), PASMCs (GSE144274) (middle), and PAAFs (GSE193776) (right). TPM - transcripts per million. **B, C**: Immunoblot analysis of indicated proteins in PASMCs and PAAFs from control and PAH subjects. Data are means±SEM; 5 subjects/group; each point represents an individual subject. *p<0.05 by Mann-Whitney U test. **D-F**: Immunoblot (**D**), proliferation (Ki67) (**E**), and apoptosis (*In Situ* Cell Death Detection Kit) (**F**) analyses of human PAH PASMCs transfected with control siRNA (-) or DEPDC1 siRNA for 48 hours. Representative images from 3 subjects/group are shown. Data are means±SEM, n=3 subjects/group, *p<0.05, *p<0.0001 by Mann-Whitney U test.

### SM-specific DLGAP5 knockdown attenuates SuHx-induced pulmonary vascular remodeling and PH in mice

To determine the contribution of PASMC-specific DLGAP5 overaccumulation to PH progression, we used an AAV-based approach. Male mice received a single tail vein injection of AAV6 encoding *EnTagln*-driven DLGAP5 shRNA or matched scrambled shRNA (Figure S3) followed by three consequent weekly injections of SU5416 and exposure to normobaric hypoxia (10% O₂). Controls were same-age same-sex normoxia-maintained mice of the same background. Hemodynamic analysis and tissue collection were performed 7 weeks after experiment initiation (Figure 5A).

**Figure 5.**
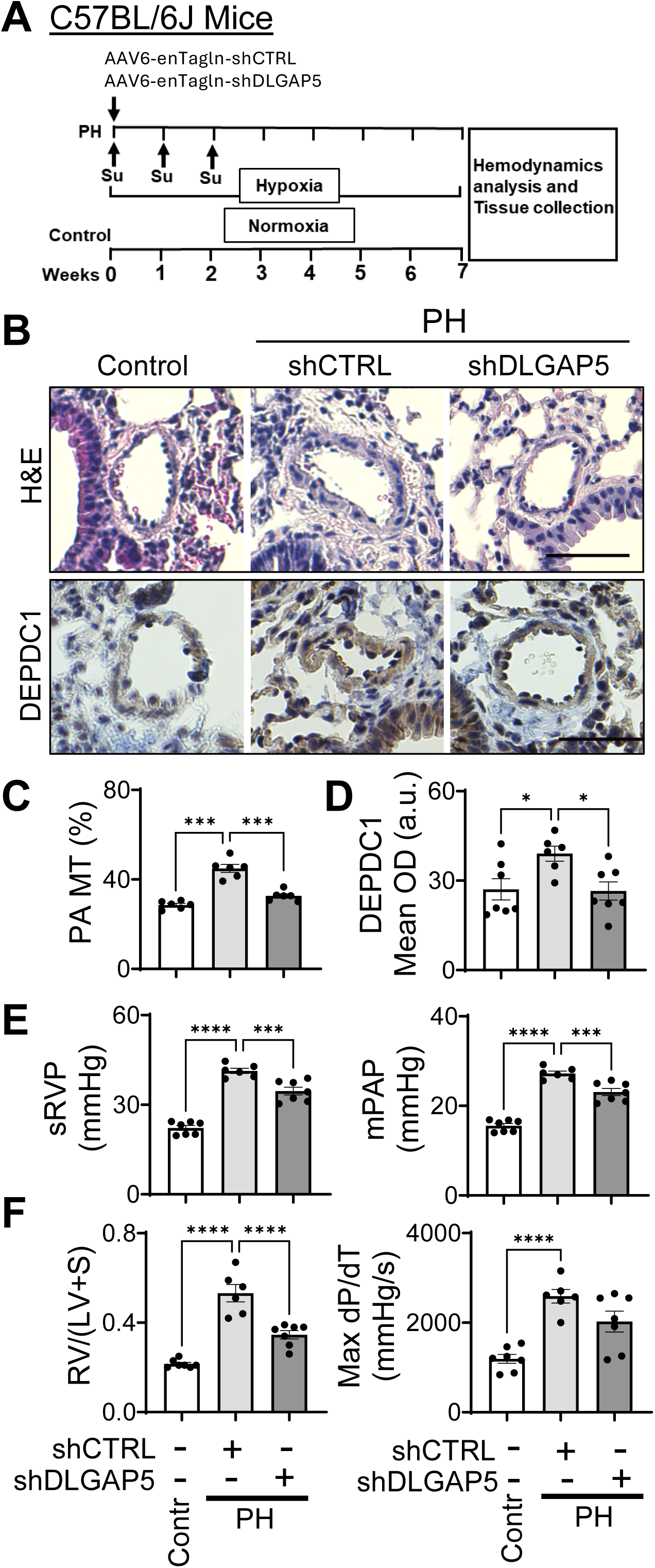
Smooth muscle-specific DLGAP5 knockdown attenuates SuHx-induced remodeling and pulmonary hypertension in mice. **A:** Male C57BL/6J mice received a single tail vein injection of AAV6 encoding *EnTagln*-driven scrambled control shRNA (AAV6-*enTagln*-shCTRL) or DLGAP5-targeting shRNA (AAV6-*enTagln*-shDLGAP5) on day 1 at a dose of 1×10¹¹ viral genomes/ mouse. Pulmonary hypertension was induced by SU5416 (20 mg/kg, sc) on days 1, 7, and 14, together with exposure to normobaric hypoxia (10% O₂). Age-matched male control mice were maintained under normoxic conditions. Seven weeks after experiment initiation, hemodynamic measurements and tissue collection were performed. **B-D:** Representative images of H&E- and anti-DEPDC1 antibodies stained lung sections (**B**), PA MT (**C**), and DRPDC1 levels (mean optical density (OD) in small muscular PAs (**D**) of mice treated as described in **A**. **E, F**: sRVP, PAP, Fulton index (RV/(LV + S), and RV max dP/dT of indicated groups are shown. Data are means±SEM; n=6–7 mice/group; *p<0.05, ***p<0.001, ****p<0.0001 by ANOVA (Dunnett’s).

SuHx-exposed AAV6-*enTagln*-shContr-injected mice developed marked PA remodeling (elevated PA MT) that was accompanied by overaccumulation of DEPDC1, increased systolic RV pressure (sRVP), mean PA pressure (mPAP), max dP/dT, and RV hypertrophy (assessed by Fulton index, RV/(LV+S) (Figure 5B-F). SMC-specific DLGAP5 knockdown reduced DEPDC1, reversed PA MT and significantly decreased sRVP, mPAP, and RV hypertrophy, but not systolic left ventricular pressure (sLVP) (Figures 5B-F, S4). There was a trend to max dP/dT improvement, which, however, did not reach statistical significance (Figure 5F). We observed no differences in body weight and heart rates among normoxia-maintained control, SuHx-exposed AAV6-*enTagln*-shContr and AAV6-*enTagln-*shDLGAP5-treated mice (Figure S4).

Collectively, these data demonstrate that SM-specific DLGAP5 knockdown attenuates SuHx-induced DEPDC1 over-accumulation in small PAs, and reduces pulmonary vascular remodeling, PH, and RV hypertrophy in mice.

## DISCUSSION

Remodeling of small muscular PAs remains a major determinant of increased pulmonary vascular resistance, elevated PA pressure and RV afterload in PAH^1,3,4,6,52–54^. This study identifies mitotic spindle-associated regulator DLGAP5 as a molecular driver of hyper-proliferation and apoptosis resistance of human PAH PASMCs and PAAFs, pulmonary vascular remodeling, and PH. Specifically, we report that DLGAP5 is overexpressed in PAH PASMCs and PAAFs, which is required for hyper-proliferation and apoptosis resistance. We also show that DLGAP5 acts through upregulating Cyclin A2 and BUBR1, and suppressing pro-apoptotic BIM, and identify DEPDC1 as a novel pro-proliferative pro-survival downstream effector of DLGAP5 in human PAH PASMCs. We further demonstrate that SMC-specific DLGAP5 knockdown attenuates pulmonary vascular remodeling, PH, and RV hypertrophy in mice with SuHx-induced PH. Together, these findings identify DLGAP5 as a disease-relevant regulator of pulmonary vascular remodeling, and DEPDC1 as its PASMC-specific downstream effector in PAH lungs.

DLGAP5 is a microtubule-associated mitotic protein that supports spindle dynamics, kinetochore capture, Ran-dependent spindle formation, and mitotic progression^30,31^. Although DLGAP5 has been linked to tumor cell proliferation, cell cycle, invasion, and cancer progression^33–35^, our findings extend its relevance to pulmonary vascular remodeling and PAH. We found that DLGAP5 overaccumulation supports increased proliferation and survival of PASMCs and PAAFs from human PAH lungs through promoting expression of cell cycle and mitosis-associated genes, and up-regulating cyclin A2 and BUBR1. Cyclin A2 promotes S phase progression and G2/M transition through cyclin-dependent kinases (CDK) 1 and 2 and participates in nuclear envelope breakdown and nuclear accumulation of Cyclin B1-CDK1 during mitotic entry^55,56^. BUBR1 is a key component of the spindle assembly checkpoint^21,22^. Coordinated reduction of mitotic gene expression, Cyclin A2 and BUBR1 in cells with DLGAP5 knockdown suggests that DLGAP5 supports pathological proliferation of pulmonary vascular cells in PAH by promoting cell cycle progression and mitotic checkpoint programs.

Previous studies have implicated the importance of mitochondrial dynamics, mitotic checkpoint signaling, and microtubule-associated processes in pulmonary vascular cell proliferation and remodeling^17,25,27–29^. Dysregulation of microtubule regulatory proteins, including BUB3, dynamin-2 (DNM2), DRP1, and glycoprotein130 (GP130), has been shown to promote hyperproliferation of pulmonary vascular cells, pulmonary vascular remodeling, and RV dysfunction in human PAH/experimental PH^17,25–27,57,58^. Although there is no known direct interactions, the possibility of indirect crosstalk between these proteins and DLGAP5 should be considered. As an example, DLGAP5-regulated cell cycle checkpoint kinases, including CDK1/Cyclin B, are shown to control DRP1-mediated mitochondrial fragmentation^59,60^, suggestive of regulatory role of DLGAP5 in cell cycle and mitochondrial dynamic signaling.

Our study identify DEPDC1 as a novel downstream effector of DLGAP5 in human PAH PASMCs. DEPDC1 acts as a regulator of cell cycle and mitosis^61,62^ and promotes cancer cell proliferation, motility and survival^61–63^. We found that DEPDC1 is overaccumulated in human PAH PASMCs and is required for increased cell proliferation and impaired apoptosis. Interestingly, PAH PAAFs did not exhibit DEPDC1 dysregulation, implying cell-specific mechanisms of DLGAP5 action in PAH pulmonary vasculature. Similar to DLGAP5, DEPDC1 affected pro-apoptotic BIM. However, it had no effect on DLGAP5-dependent BUBR1 and Cyclin A2, indicative of multi-faceted DLGAP5-directed downstream signaling that requires further investigation.

Our findings have preclinical relevance to the evolving therapeutic landscape of PAH. Majority of FDA-approved PAH therapies target three major vasoactive pathways, including endothelin, nitric oxide, and prostacyclin signaling. These therapeutics have limited effect on pulmonary vascular remodeling and don’t stop disease progression^1,9,10^. Recent approval of anti-proliferative Sotatercept have reinforced the importance of targeting mechanisms that directly regulate pulmonary vascular cell growth, survival, and remodeling^10^. In sync with our *in vitro* data, identifying DLGAP5 as an important mitotic regulator that contributes to hyperproliferation of pulmonary vascular cells, our *in vivo* findings support pathophysiological importance of DLGAP5 in experimental pulmonary vascular remodeling and PH. In the SuHx mouse model, SMC-specific DLGAP5 knockout down-regulated DEPDC1, reversed PA remodeling, lowered PA pressure, and attenuated RV hypertrophy. It should be noted, however, that SMC-specific DLGAP5 silencing did not completely reverse sRVP and mPAP. This could be explained, at list in part, by selective targeting of SMC, which didn’t improve DLGAP5-specific PAAF pathobiology. It is possible that combined targeting of PASMC and PAAF is needed to completely reverse disease.

Several limitations should be acknowledged. First, the human tissue- and primary cell-based experiments were performed using a limited number of specimens from patients with PAH and non-diseased donors. PAH is a rare disease, and access to well-characterized human lung tissues and early-passage primary pulmonary vascular cells is limited. However, the involvement of DLGAP5 in PAH vascular remodeling is supported by complementary analyses of human PAH transcriptomic datasets, patient-derived PASMCs and PAAFs, siRNA-mediated loss-of-function studies, and smooth muscle-targeted AAV6-mediated DLGAP5 knockdown in the SuHx mouse model. Second, although our data identify DEPDC1 as a downstream effector of DLGAP5, the precise mechanism by which DLGAP5 regulates DEPDC1 protein levels and nuclear accumulation remains to be defined. Third, the *in vivo* studies were performed in male mice; therefore, validation in female animals, additional experimental models, and conditional genetic approaches would further strengthen the generalizability of these findings.

In summary, our study provides new evidence that DLGAP5 supports hyperproliferative, apoptosis-resistant phenotype of pulmonary vascular cells in PAH, identify DEPDC1 as a novel downstream effector of DLGAP5 and demonstrate that DLGAP5 targeting attenuates pulmonary vascular remodeling and overall PH. Our findings suggest that DLGAP5 may represent a potential therapeutic target to resolve pulmonary vascular remodeling and reverse PAH.

## PERSPECTIVES

Despite therapeutic advances, the mechanisms of pulmonary vascular remodeling remain incompletely defined. Our findings position DLGAP5 and its downstream effectors DEPDC1, BUBR1, and Cyclin A2 as contributors to PAH pulmonary vascular remodeling and support further preclinical investigation of DLGAP5-regulated pathways. Future work is needed to delineate the relationship between DLGAP5 and DEPDC1, establish the cell type- and sex-specificity of these effects, and validate their relevance across experimental models and human PAH subtypes. Determining whether DLGAP5-associated mitotic programs can be selectively modulated will be important for assessing DLGAP5 translational potential.

## NOVERLTY AND RELEVANCE

### What Is New?

- DLGAP5, a mitotic spindle-associated regulator, is elevated in remodeled pulmonary arteries, pulmonary artery smooth muscle cells and adventitial fibroblasts from patients with PAH.
- DEPDC1 is identified as a novel downstream effector of DLGAP5.
- Smooth muscle-specific DLGAP5 silencing attenuates pulmonary vascular remodeling, PH, and right ventricular hypertrophy in mice with experimental pulmonary hypertension.

### What Is Relevant?

Pathological pulmonary vascular cell hyper-proliferation and apoptosis resistance are central drivers of pulmonary vascular remodeling in PAH. Our findings identify DLGAP5 as molecular driver of the hyperproliferative, apoptosis-resistant phenotype of PAH pulmonary vascular cells, and pulmonary vascular remodeling and PH in mice, extending our knowledge about mitotic regulatory pathways contributing to pulmonary vascular remodeling.

### Clinical/Pathophysiological Implications?

DLGAP5 could represent new remodeling-focused potential molecular target to treat PAH.

