## Supplementary material for "DLGAP5 promotes vascular cell proliferation and remodeling in pulmonary arterial hypertension": Online supplement

### Materials and Methods

#### Human tissues and primary pulmonary vascular cell culture

The human lung tissues and early-passage (3-8) human pulmonary arterial smooth muscle cells (PASMCs), pulmonary arterial endothelial cells (PAECs), and pulmonary arterial adventitial fibroblasts (PAAFs) from small muscular PAs (<1.5 mm outer diameter) of non-diseased donors and patients with PAH were provided by the Pulmonary Hypertension Breakthrough Initiative (PHBI), the University of California, San Francisco lung transplant program, and the University of California, Davis Lung Center Pulmonary Vascular Disease Program biobank (tissue acquired from the University of Pittsburgh Medical Center Lung Transplant program and Pulmonary Division Tissue Donation program, Institutional Review Board (IRB) approvals #PRO14010265, #PRO13050218 and Committee for Oversight of Research Involving Decedents approvals #451, #458), and the University of California San Francisco Lung Transplant Program (IRB protocol # 13-10738).

Pulmonary vascular cells were isolated and maintained using the PHBI approved protocols[1, 2]. Precapillary PAs were microdissected from surrounding lung tissue, and adherent parenchyma was removed. For PASMC and PAAF isolation, non-medial and non-adventitial layers, respectively, were mechanically removed, remaining tissue was minced and plated in lineage-specific complete growth media: Smooth Muscle Cell Growth Medium 2 (PromoCell, Heidelberg, Germany; Cat. No. C-22162) for PASMCs, Fibroblast Growth Medium 2 (PromoCell; Cat. No. C-23120) for PAAFs. PAECs were isolated using enzymathic digestion and maintained in Endothelial Cell Growth Medium MV (PromoCell; Cat. No. C-22120). All cultural media were supplemented with Antibiotic-Antimycotic (Thermo Fisher Scientific, Waltham, MA; Cat. No. 15240062).

Experiments were performed using matched passages from at least three different subjects. Before experiments, cells were maintained in basal medium supplemented with 0.1% bovine serum albumin (BSA) (PASMCs and PAAFs) or 0.5% fetal bovine serum (PAECs) for 48 hours unless otherwise indicated.

#### Animals

All animal procedures were performed under the protocols approved by the Animal Care and Use Committee of the University of California, Davis. All experimental groups were assigned by randomization; hemodynamic and morphologic analyses were performed in a blinded manner. 12-week-old male C57BL/6J mice were purchased from the Jackson Laboratory (Bar Harbor, ME, Cat. No. 000664). On day 1, mice received a single tail vein injection of AAV6 ($1 \times 10^{11}$ viral genomes/mouse) (VectorBuilder Inc, Chicago, IL) encoding either shDLGAP5 or scrambled shRNA under the control of the SM22α promoter. The construct pAAV[3miR30]-*enTagln*-EGFP-mDlgap5(shRNAs)-WPRE

[VectorBuilder construct ID AAV6SP(VB241001-1545nrb)] and the corresponding scrambled-control construct was pAAV[3miR30]-*enTagln*-EGFP-scramble(shRNAs)-WPRE [VectorBuilder construct ID AAV6SP(VB250618-1675urv)] purchased form VectorBuilder. The AAV6-shDLGAP5 vector contained three miR30-based shRNAs targeting mouse *Dlgap5*. The target sequences were 5'-ACATTCAGTGCTCGCCTTTAAT-3', 5'-AGCGAAACAGACACTTCGGTTT-3', and 5'-TCTTCTGAACGTCGTTCTCAAA-3'. On days 1, 7, and 14, mice received SU5416 (20 mg/kg, s.c.) and exposed to normobaric hypoxia (10% $O_2$) for 7 weeks. Age-matched male mice maintained under normoxic conditions were used as controls. Seven weeks after experiment initiation, blinded hemodynamic measurements were performed[1-4]. Mice were anesthetized with isoflurane (Cat. No.803250, McKesson Medical Surgical Inc, Richmond, VA; induction at 2%, maintenance at 1–2%). RV and LV pressure recordings were obtained using a pressure-volume catheter (Cat. No. FTH-1212B-3518, Transonic Scisense Inc, Ithaca, NY). Data were recorded and analyzed using the LabChart Pro system (ADInstruments Inc, Colorado Springs, CO). Mean PA pressure was calculated as 0.61 x sRVP + 2 (in mmHg)[5]. Fulton index was calculated as [RV/(LV+S)] weight ratio[3]. For morphological and immunohistochemical analyses, lungs were perfused with phosphate-buffered saline (PBS) through the PA, fixed in 4% paraformaldehyde, and paraffin embedded. Lung sections (5 μm) were stained with hematoxylin and eosin (H&E) or subjected to immunohistochemical staining[1]. Images were acquired from randomly selected small pulmonary arteries (≤150 μm outer diameter) from at least six animals per group, with a minimum of 10 PAs per animal. PA medial thickness (PA MT) was calculated using the VMI calculator[6].

### Immunohistochemical analysis

Immunohistochemical analysis was performed using methods as previously described[1,3,7]. Briefly, human lung tissues were fixed in 4% paraformaldehyde in PBS, paraffin-embedded, and sectioned (5μm thickness). Tissue sections were processed for immunofluorescent immunohistochemical (IHC) staining according to protocols described in[1,3]. Paraffin-embedded lung tissue sections were deparaffinized in xylene and rehydrated through a series of graded ethanol to distilled water. Tissues were then permeabilized by Triton X-100 (0.1 %) in PBS. For immunofluorescent staining, heat-induced antigen retrieval was performed using citrate buffer (Diagnostic Biosystems, Pleasanton, CA, Cat. No. NC2049147) by manufacturer's instructions. Then, sections were blocked with 2% BSA and then incubated with respective primary and secondary antibodies. Images were acquired under identical settings across the experimental groups. For 3,3′-diaminobenzidine (DAB) staining, heat-induced antigen retrieval was performed using Epitope Retrieval Buffer (IHC World, Ellicott City, MD, Cat. No. IW-1100-1L) according to manufacturer's protocol. After cooling to room temperature, sections were washed with PBS, and endogenous peroxidase activity was quenched with 3% hydrogen peroxide. Sections were blocked with the normal serum supplied with the VECTASTAIN ABC-HRP Kit (Vector Laboratories, Newark, CA; Cat. No. PK-4001) and incubated overnight at 4°C with primary and secondary antibodies. After washing, sections were incubated with the appropriate biotinylated secondary antibody, followed by the avidin-biotin-horseradish peroxidase complex prepared according to the manufacturer's instructions. Signals were developed using DAB, and the reaction was stopped at the desired brown signal intensity. Sections were counterstained with hematoxylin, dehydrated through graded ethanol solutions, cleared in xylene, and cover-slipped using a xylene-based permanent mounting medium. Images were acquired using a BZ-X810 All-in-One fluorescence microscope under identical settings across the experimental groups. For quantification, PA walls were manually selected in ImageJ, and the lumen was excluded from the region of interest. DAB signal was separated using Color Deconvolution2 with the H-DAB vector (8-bit transmittance output), and a fixed threshold of 0-125 was applied uniformly across all images. DEPDC1-positive staining was expressed as mean optical density (OD). Ten PAs per animal were analyzed, and values were averaged to obtain a single value for each animal for statistical analysis.

Primary antibodies used for immunohistochemical analysis included rabbit polyclonal anti-DLGAP5 (ABclonal, Woburn, MA; Cat. No. A2197; 1:80), rabbit polyclonal anti-DEPDC1 primary antibody (Bioss, Cat. No. bs-6525R; 1:300), FITC-conjugated mouse monoclonal anti–α-smooth muscle actin (α-SMA; clone 1A4; MilliporeSigma, St. Louis, MO; Cat. No. F3777; 1:150). Alexa Fluor 594–conjugated chicken anti-rabbit IgG (H+L) (Thermo Fisher Scientific Cat. No. A-21442; 1:500) secondary antibody was used for signal detection. The FITC-conjugated α-SMA antibody was detected directly without a secondary antibody. Nuclei were counterstained with 4′,6-diamidino-2-phenylindole (DAPI; Thermo Fisher Scientific; Cat. No. D21490). Fluorescence images were acquired using a BZ-X810 All-in-One fluorescence microscope.

**Immunoblot analysis** was performed as previously described[1, 2]. Cells were lysed in buffer A containing 40 mmol/L HEPES, 120 mmol/L NaCl, 10 mmol/L sodium pyrophosphate, 10 mmol/L β-glycerophosphate, 50 mmol/L NaF, and 1 mmol/L EDTA (Thermo Fisher Scientific; Cat. No. 15575020), supplemented with sodium orthovanadate (1.5 mmol/L; MilliporeSigma; Cat. No. 567540) and protease inhibitor cocktail (Thermo Fisher Scientific; Cat. No. PI78415). Protein concentration was determined using the bicinchoninic acid (BCA) assay.

Protein lysates were mixed with NuPAGE LDS sample buffer (Thermo Fisher Scientific; Cat. No. NP0007) and NuPAGE sample reducing agent (Thermo Fisher Scientific; Cat. No. NP0009). Samples were heated at 70°C for 10 minutes, run in 4%–12% NuPAGE Bis-Tris gels (Thermo Fisher Scientific; Cat. No. NP0323BOX), and transferred to nitrocellulose membranes. The membranes were blocked in 5% nonfat dry milk, incubated overnight at 4°C with primary antibodies, and later with horseradish peroxidase (HRP)–conjugated secondary antibodies. The signal was detected by enhanced chemiluminescence reagent and visualized by exposure to HyBlot CL autoradiography film (Thomas Scientific, Swedesboro, NJ; Cat. No. 1156P37).

Primary antibodies were mouse monoclonal anti-DLGAP5/HURP (clone E-7; Santa Cruz Biotechnology, Dallas, TX; Cat. No. sc-377004; 1:500), rabbit polyclonal anti–α/β-tubulin (Cell Signaling Technology, Danvers, MA; Cat. No. 2148; 1:2000), rabbit monoclonal anti-BIM (clone C34C5; Cell Signaling Technology; Cat. No. 2933; 1:1000), rabbit monoclonal anti-Cyclin A2 (clone E1D9T; Cell Signaling Technology; Cat. No. 91500; 1:1000), rabbit polyclonal anti-BubR1 (Bioss; Cat. No. bs-5726R; 1:1000), and rabbit polyclonal anti-DEPDC1 (Invitrogen, Carlsbad, CA; Cat. No. PA5-70206; 1:1000). Secondary antibodies were HRP-conjugated anti-rabbit IgG (Cell Signaling Technology; Cat. No. 7074; 1:1000–1:2500) and HRP-conjugated anti-mouse IgG (Thermo Fisher Scientific; Cat. No. A16072; 1:2000).

**Cell Transfection** was performed as previously described[1, 4]. Briefly, cells were transfected with ON-TARGETplus small interfering RNAs (siRNAs) targeting DLGAP5 (Dharmacon, Lafayette, CO; Cat. No. L-016846-00-0020), DEPDC1 (Dharmacon, Cat. No. L-021125-00-0010) or a nontargeting control siRNA #1 (Dharmacon; Cat. No. D-001810-01-20) using DharmaFECT 1 transfection reagent (Horizon Discovery Biosciences, Lafayette, CO; Cat. No. T-2001-03). Transfections were carried out according to the manufacturer's protocol, cells were analyzed 48 hours after transfection.

**RNA extraction and bulk RNA sequencing**

PASMCs from three patients with PAH were transfected with siDLGAP5 or nontargeting control siRNA (siCTRL) for 48 hours. Cells were lysed in TRIzol reagent and submitted to Genewiz® (Azenta Life Sciences, South Plainfield, NJ) for total RNA extraction and quality assessment. Stranded RNA-seq libraries were prepared with poly(A) selection and sequenced on the Illumina NovaSeq platform, generating 2×150-bp paired-end reads at a depth of approximately 50 million reads per sample. Library construction, sequencing quality control, read trimming and alignment, transcript quantification, normalization, and differential gene-expression analysis were performed by Genewiz®. Differential-expression results comparing siDLGAP5- and siCTRL-treated PASMCs were used for downstream analyses.

**Public transcriptomic data collection and analysis**

Publicly available human PAH transcriptomic datasets were obtained from the Gene Expression Omnibus. GSE113439[8] contains microarray expression profiles of PAH and control lungs, whereas GSE144274[9], GSE193776[10], and GSE126262[11] contain RNA-sequencing profiles of PASMCs, PAAFs, and PAECs, respectively, from patients with PAH and nondiseased controls. Processed expression data normalized by the original studies were used, including robust multiarray average-normalized values for the microarray dataset and normalized transcript-abundance values for the RNA-sequencing datasets[12]. DLGAP5 and DEPDC1 expression levels were compared between PAH and control samples using the Mann-Whitney U test.

**STRING protein–protein network analysis**

Downregulated DEGs identified by bulk RNA sequencing of human PAH PASMCs following DLGAP5 silencing were analyzed using the STRING database (version 12.5; accessed September, 2025). Homo sapiens was selected as the reference organism, and protein-protein interactions were retrieved using a confidence score of 0.95. The network was then visualized using STRING version 12.5. Nodes represent proteins encoded by downregulated genes, and edges represent known or predicted functional protein associations.

**Proliferation and apoptosis assays**

Cell proliferation was assessed by immunocytochemical detection of Ki-67 as previously described[1, 3, 4]. Briefly, cells were washed with PBS, fixed in 4% paraformaldehyde in PBS, permeabilized with Triton X-100 in PBS, and blocked in 2% BSA in PBS. Cells were then incubated overnight at 4°C with a Ki-67 (D3B5) Rabbit mAb primary antibody (Cell Signaling Technology; Cat. No. 9129; 1:400) diluted in blocking solution. Following PBS washes, cells were incubated with Chicken anti-Rabbit IgG (H+L) Cross-Adsorbed Secondary Antibody, Alexa Fluor™ 594 (Thermo Fisher Scientific; Cat. No. A-21442; 1:500). Nuclear counterstaining was performed with DAPI (Thermo Fisher Scientific, Cat. No. D21490).

Apoptotic cell death was evaluated using the In Situ Cell Death Detection Kit (TUNEL), Fluorescein (Millipore Sigma/Roche; Cat. No. 11684795910), according to the manufacturer's instructions. Nuclei were counterstained with DAPI. Fluorescent images were acquired using a BZ-X810 All-in-One fluorescence microscope.

. TUNEL-positive and Ki67-positive cells were counted manually in a blinded manner using open source graphics editor GIMP 2.10. Cell proliferation was calculated as the percentage of Ki67-positive cells relative to the total number of cells, and apoptosis was quantified as the percentage of TUNEL-positive cells relative to total number of nuclei. A minimum of 200 cells per condition were analyzed in each experiment.

**Immunocytochemical analysis and quantification of nuclear DEPDC1 mean fluorescence intensity** Immunocytochemical analysis was performed as previously described[4]. Briefly, cells were fixed in paraformaldehyde (4%), permeabilized with Triton X-100 (0.1%) in PBS, and blocked with 2% BSA. Cells were incubated with DEPDC1 (Bioss, Cat. No. bs-6525R; 1:200) primary antibody and Alexa Fluor 594–conjugated chicken anti-rabbit IgG (H+L) (Thermo Fisher Scientific, Cat No. A-21442; 1:500) secondary antibody. Nuclei were counterstained with DAPI. Fluorescence images were acquired using a BZ-X810 All-in-One fluorescence microscope.

Nuclear DEPDC1 fluorescence intensity was quantified using ImageJ. DAPI channel images were used to define nuclear regions of interest (ROIs). Briefly, DAPI images were converted to 8-bit grayscale, converted to binary masks, and processed using Fill Holes and Watershed to separate adjacent nuclei. Nuclear ROIs were generated using Analyze Particles and saved in the ROI Manager. The DAPI-derived nuclear ROIs were then applied to the corresponding DEPDC1 fluorescence channel from the same image field. DEPDC1 mean fluorescence intensity (MFI) was measured within each nuclear ROI using identical image acquisition settings for all the samples. Background fluorescence was measured from cell-free regions of each image and subtracted from nuclear DEPDC1 MFI. Data are presented as background-corrected nuclear DEPDC1 MFI in arbitrary units (a.u.). For quantification of nuclear DEPDC1-positive cells, a fixed background-corrected nuclear DEPDC1 MFI cutoff of >11,000 arbitrary units was applied to define DEPDC1-positive nuclei. This cutoff was applied uniformly across all the samples.

**Statistical analysis**

Statistical analyses were performed using StatView and GraphPad Prism. For datasets with $n<6$ per group, comparisons between two groups were performed using the Mann-Whitney U test. Normally distributed datasets with $n\geq6$ per group were analyzed using Student's t-test for two-group comparisons or one-way ANOVA with Dunnett's post hoc correction for multiple-group comparisons. Statistical significance was defined as $p<0.05$.

**Supplemental Tables and Figures**

**Table S1. Human subjects' characteristics**

| Diagnosis | Gender | Age |
|---|---|---|
| Non-Diseased | F | 36 |
| Non-Diseased | M | 47 |
| Non-Diseased | F | 50 |
| Non-Diseased | M | 19 |
| Non-Diseased | F | 33 |
| Non-Diseased | M | 36 |
| Non-Diseased | M | 38 |
| Non-Diseased | F | 48 |
| Non-Diseased | M | 40 |
| Non-Diseased | F | 36 |
| Non-Diseased | F | 57 |
| Non-Diseased | F | 53 |
| Non-Diseased | M | 25 |
| Non-Diseased | M | 47 |
| IPAH | F | 40 |
| IPAH | F | 43 |
| PAH Scleroderma | F | 53 |
| PAH Scleroderma | F | 62 |
| IPAH | F | 50 |
| PAH | F | 36 |
| IPAH | M | 44 |
| IPAH | M | 21 |
| IPAH | M | 73 |
| IPAH | M | 53 |
| IPAH | F | 27 |
| IPAH | M | 16 |
| IPAH | F | 16 |
| IPAH | M | 53 |

F - Female; M - Male; IPAH – Idiopathic pulmonary arterial hypertension

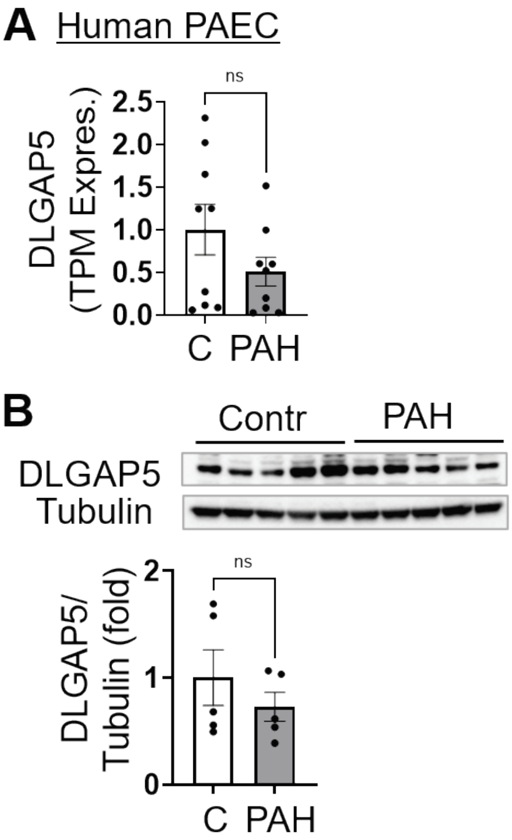


**Figure S1. A:** DLGAP5 expression in human pulmonary artery endothelial cells; GSE126262 dataset, n=9 controls, n=10 IPAH subjects. TPM - relative transcripts per million. **B**: Immunoblot analysis of early-passage human PAECs from control and PAH subjects to detect the indicated proteins. n=5 subjects/group. Data are means±SE. ns - non-significant by Mann-Whitney U test.

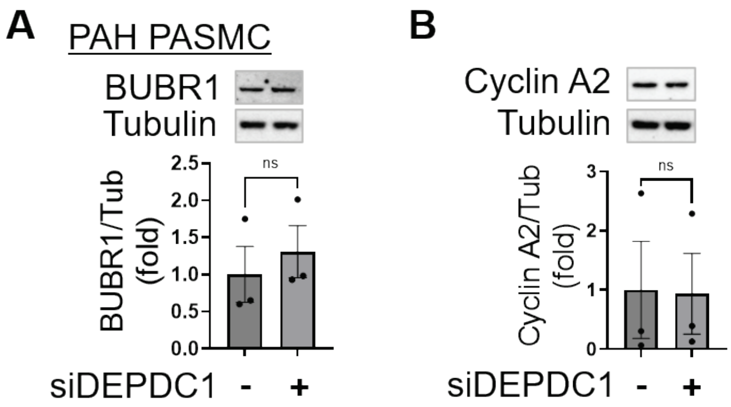


**Figure S2**. Immunoblot analysis of PAH PASMCs transfected with scrambled control siRNA or DEPDC1 siRNA for 48 hours to detect indicated proteins. Data are means±SEM. n=3 subjects per group; ns- non-significant by Mann-Whitney U test

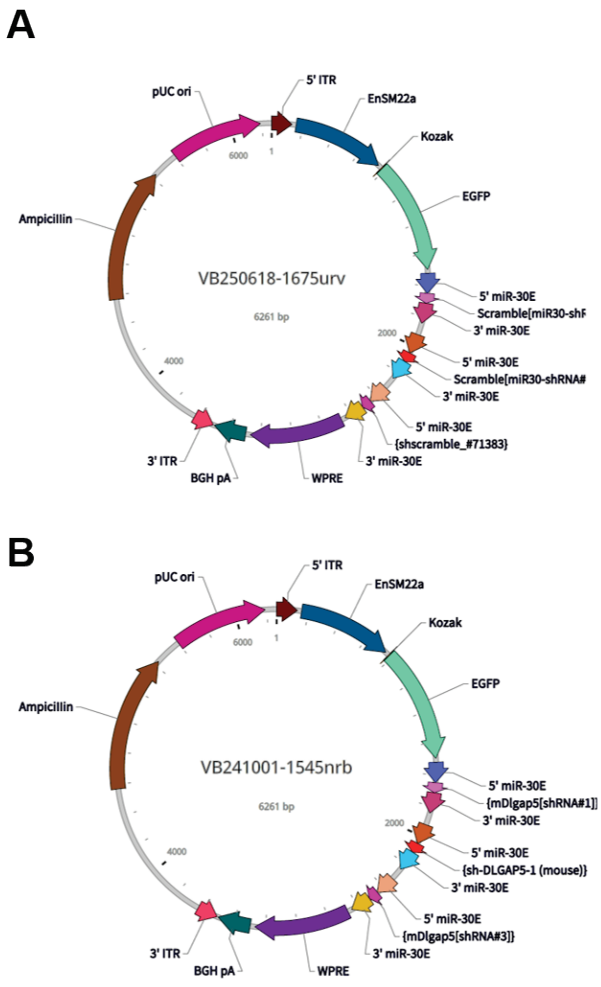


**Figure S3**. Design of mammalian Triple miR30-shRNA Knockdown AAV Vectors. **A**: Scramble shRNA (Vector Id VB250618-1675urv). **B**: shDLGAP5 (Vector ID: VB241001-1545nrb).

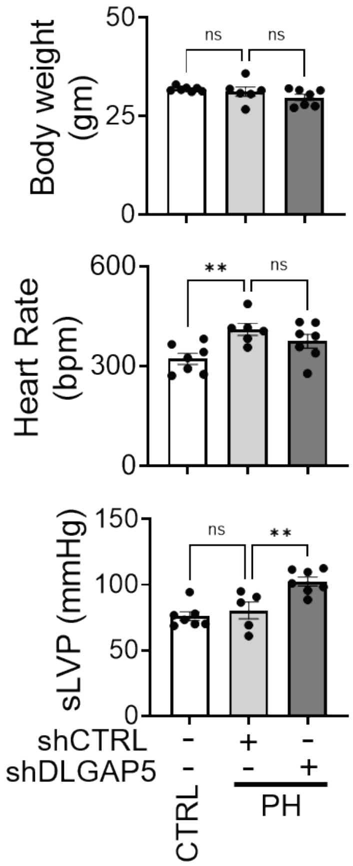


**Figure S4**. Endpoint body weight, heart rate, and systolic LV pressure (sLVP) were measured in control mice and mice with SU5416/hypoxia-induced pulmonary hypertension (PH) treated with AAV6-shCTRL or AAV6-shDLGAP5. Data are means±SEM; n=6-7 mice per group. **p<0.01, ns - not significant by one-way ANOVA (Dunnett's posthoc correction).
